# Predicting stop codon reassignment improves functional annotation of bacteriophages

**DOI:** 10.1101/2023.12.19.572299

**Authors:** Ryan Cook, Andrea Telatin, George Bouras, Antonio Pedro Camargo, Martin Larralde, Robert A. Edwards, Evelien M. Adriaenssens

## Abstract

The majority of bacteriophage diversity remains uncharacterised, and new intriguing mechanisms of their biology are being continually described. Members of some phage lineages, such as the *Crassvirales*, repurpose stop codons to encode an amino acid by using alternate genetic codes. Here, we investigated the prevalence of stop codon reassignment in phage genomes and subsequent impacts on functional annotation. We predicted 76 genomes within INPHARED and 712 vOTUs from the Unified Human Gut Virome catalogue (UHGV) that repurpose a stop codon to encode an amino acid. We re-annotated these sequences with modified versions of Pharokka and Prokka, called Pharokka-gv and Prokkagv, to automatically predict stop codon reassignment prior to annotation. Both tools significantly improved the quality of annotations, with Pharokka-gv performing best. For sequences predicted to repurpose TAG to glutamine (translation table 15), Pharokka-gv increased the median gene length (median of per genome medians) from 287 to 481 bp for UHGV sequences (67.8% increase) and from 318 to 550 bp for INPHARED sequences (72.9% increase). The re-annotation increased mean coding density from 66.8% to 90.0%, and from 69.0% to 89.8% for UHGV and INPHARED sequences. Furthermore, the proportion of genes that could be assigned functional annotation increased, including an increase in the number of major capsid proteins that could be identified. We propose that automatic prediction of stop codon reassignment before annotation is beneficial to downstream viral genomic and metagenomic analyses.

## Main Body

Bacteriophages, hereafter phages, are increasingly recognised as a vital component of microbial communities in all environments where they have been studied in detail. Phages are known to drive bacterial evolution and community composition through predator-prey dynamics and their potential as agents of horizontal gene transfer. The use of viral metagenomics, or viromics, has massively expanded our understanding of global viral diversity and shed light on the ecological roles that phages play.

Much of the study into viral communities has been conducted on the human gut. Here, viromics has uncovered ecologically important viruses that are difficult to bring into culture using standard laboratory techniques^1^, shown potential roles of viruses in disease states^2^, and allowed for the recovery of enormous phage genomes larger than any brought into culture^3^. As the majority of phage diversity remains uncharacterised, new and enigmatic diversification mechanisms are being described continually, including the potential use of alternative translation tables.

Lineage-specific stop codon reassignment has been described previously in bacteriophages^4,5^, whereby a stop codon is repurposed to encode an amino acid. Notably, annotations of Lak “megaphages” assembled from metagenomes were observed to exhibit unusually low coding density (∼70%) when genes are predicted using the standard bacterial, archaeal and plant plastid genetic code (translation table 11)^3^, much lower than the value observed for most cultured phages of ∼90%^6^. The Lak megaphages were predicted to repurpose the TAG stop codon into an as-of-yet unknown amino acid^3^. More recently, uncultured members of *Crassvirales* have been predicted to repurpose TAG to glutamine (translation table 15), and TGA to tryptophan (translation table 4)^5^, and since then the use of translation table 15 has been experimentally validated in two phages belonging to *Crassvirales*^7^. As this feature may be widespread in human gut viruses, we trained a fork of Prodigal^8^, named prodigal-gv, to predict stop codon reassignment in phages^9^ and implemented in the pyrodigal-gv library to provide efficient Cython bindings to Prodigal-gv with pyrodigal^10^. Additionally, the virus discovery tool geNomad incorporates pyrodigal-gv to predict stop codon reassignment for viral sequences identified in metagenomes and viromes^9^. However, the detection of translation table 15 still has limited support in many tools, and the impacts of stop codon reassignment are rarely considered in viral genomics and metagenomics.

To assess the extent of stop codon reassignment in studied phage genomes and the impacts on functional annotation, we extracted phage genomes from INPHARED^6^ and predicted those using alternative stop codons. We also added high-quality and complete vOTUs from the Unified Human Gut Virome Catalog (UHGV; https://github.com/snayfach/UHGV) predicted to use alternative codons. The viral genomes were re-annotated using modified versions of the commonly used annotation pipelines Prokka^11^, and Pharokka^12^ implementing prodigal-gv/pyrodigal-gv for gene prediction (Supplementary Methods). Hereafter, the modified versions are referred to Prokka-gv and Pharokka-gv.

From INPHARED, 49 genomes (0.24%) were predicted to use translation table 15, and 27 (0.13%) were predicted to use translation table 4. From the UHGV, 666 vOTUs (1.2%) were predicted to use translation table 15 and 46 (0.08%) were predicted to use translation table 4. These genomes and vOTUs were not constrained to one particular clade of viruses, being predicted to occur on both dsDNA viruses of the realm *Duplodnaviria* and ssDNA viruses of the realm *Monodnaviria*, suggesting it is a phenomenon that has arisen on at least two occasions (Supplementary Table 1). The lower frequency of these genomes in cultured isolates (INPHARED) versus human viromes (UHGV) may be due to culturing and sequencing biases, perhaps including modifications to DNA that are known to be recalcitrant to sequencing.

Although the mechanism for stop codon reassignment in phages is not fully understood, suppressor tRNAs are suggested to play a role^4,13^. Consistent with previous findings, we found 375/715 (52.4%) phages predicted to use translation table 15 encoded at least one suppressor tRNA corresponding to the amber stop codon (Sup-CTA tRNA), and 11/73 (15.1%) of those predicted to use translation table 4 encoded at least one suppressor tRNA corresponding to the opal stop codon (Sup-TCA tRNA)^4,13,14^. Although fewer of those predicted to use translation table 4 encoded the relevant suppressor tRNA, 22/27 (81%) of the INPHARED phages predicted to use translation table 4 were viruses of *Mycoplasma* or *Spiroplasma*. As *Mycoplasma* and *Sprioplasma* are known to use translation table 4, many of the viruses predicted to use translation table 4 may be simply using the same translation table as their host.

Prediction of stop codon reassignment led to improved annotations for both Prokka and Pharokka, although the extent of this varied with the two datasets, translation tables, and annotation pipelines tested. As Pharokka-gv outperformed Prokka-gv on all metrics tested, only Pharokka-gv is discussed further, and the equivalent results for Prokka-gv can be found in Supplementary Results.

The largest differences were observed for sequences predicted to use translation table 15, for which Pharokka-gv increased the median gene length (median of per genome medians) from 287 to 481 bp for UHGV sequences (67.8% increase) and from 318 to 550 bp for INPHARED sequences (72.9% increase; Figure 1A). This was also reflected in an increase of median coding capacity from 66.8% to 90.0% for UHGV, and 69.0% to 89.8% for INPHARED (Figure 1B). Overall, these improved gene calls led to an increased gene length, and a reduction in the number of predicted genes per kb and the number of genes that could not be assigned functional annotations (Supplementary Figure 2; Supplementary Table 2). As it is commonly used as a phylogenetic marker for bacteriophages, we investigated how commonly the major capsid protein (MCP) could be identified with and without predicted stop codon reassignment^15^. For those viruses we predicted to use translation table 15, annotation using the default translation table 11 only resulted in the MCP being identified in 407/715 (56.9%) of the genomes. In contrast, using translation table 15 with Pharokka-gv, we could identify the MCP in 475/715 (66.4%).

**Figure 1.**
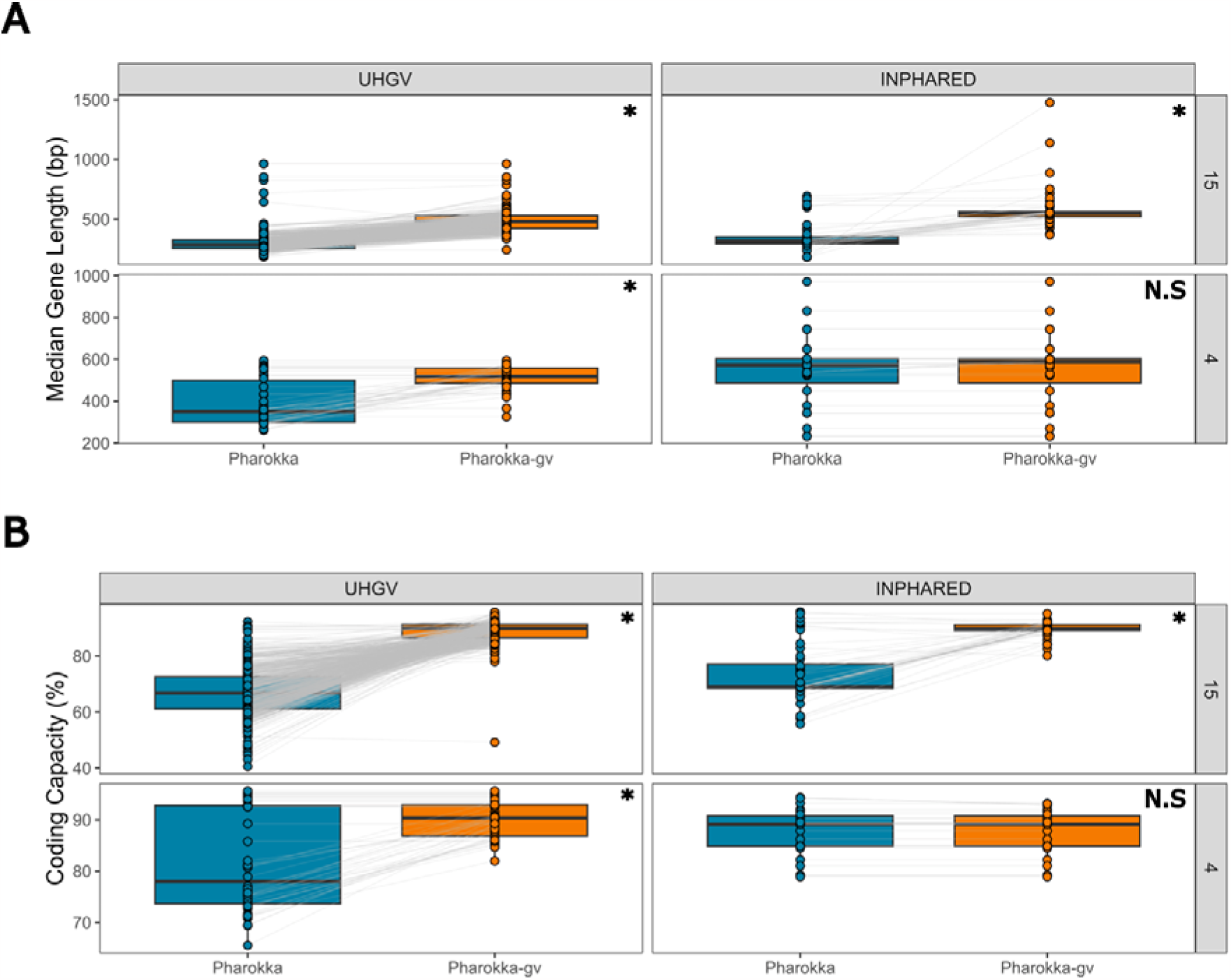
Re-annotating with predicted stop codon reassignment increases the quality of annotations. Comparison of (**A**) median predicted gene length (bp) and (**B**) coding capacity (%) for INPHARED genomes and UHGV vOTUs annotated with Pharokka (translation table 11 only) and Pharokka-gv (prediction of stop codon reassignment), grouped by dataset and predicted stop codon reassignment. Asterisk indicates significance at P ≤ 10e-10 with P determined by a simple T test and adjusted with the Benjamini-Hochberg procedure.

When investigating the sequences for which translation table 4 was predicted to be optimal, a substantial increase was also observed for UHGV sequences, with Pharokka-gv increasing median gene length (median of per genome medians) from 350 to 518 bp (a 48.0% increase in length; Figure 1A), resulting in an increase of coding capacity from 78.0% to 90.4% (Figure 1B). However, the same was not observed for the 27 INPHARED genomes predicted to use translation table 4. Reannotation resulted in a modest increase in median gene length (median of per genome medians) from 573 to 588 bp (a 2.6% increase in length; Figure 1A). Median coding capacity was not increased, with both Pharokka and Pharokka-gv obtaining 89.1% (Figure 1B). As the median gene length and coding capacity for INPHARED sequences predicted to use translation table 4 are in line with expected values, their prediction may be a false positive. Reassuringly, the prediction of translation table 4 has not hindered the quality of annotations where it may be a false positive.

The analysis of viral (meta)genomes relies on accurate protein predictions, with predicted ORFs being used in common analyses, including (pro)phage prediction, functional annotation, and phylogenetic analyses. The clear differences in protein predictions with/without predicted stop codon reassignment will likely have downstream impacts upon these analyses. However, this phenomenon is not yet widely considered in viral (meta)genomics. We have demonstrated the impacts of stop codon reassignment in the functional annotation of phages, and provide tools for the automatic prediction and annotation of viral genomes that repurpose stop codons. Our analysis highlights the need for accurate viral ORF prediction, and further experimental validation to elucidate the mechanisms of stop codon reassignment.

## Data Availability

The genomes used in this analysis are from two publicly available datasets; INPHARED (https://github.com/RyanCook94/inphared) and the Unified Human Gut Virome (UHGV; https://github.com/snayfach/UHGV). The details of included sequences are shown in Supplementary Table 1. The code for Prokka-gv is available on GitHub (https://github.com/telatin/metaprokka). The code for Pharokka is available on GitHub (https://github.com/gbouras13/pharokka). The code for Prodigal-gv is available on GitHub (https://github.com/apcamargo/prodigal-gv). The code for Pyrodigal-gv is available on GitHub (https://github.com/althonos/pyrodigal-gv).

## Supplementary Methods

### Datasets

A multifasta file of phage genomes was downloaded from INPHARED (https://github.com/RyanCook94/inphared; September 2023)^6^. Stop codon reassignment of INPHARED genomes was predicted using Prodigal-gv v2.11.0 (https://github.com/apcamargo/prodigal-gv), a fork of Prodigal written to improve viral gene calling^8^. Those predicted to use translation table 4 or 15 were retained for downstream analysis.

The Unified Human Gut Virome Catalog (UHGV) was filtered for high quality and complete vOTUs deemed to be a “high confidence” virus and predicted to use either translation table 4 or 15 (https://github.com/snayfach/UHGV). Stop codon reassignment had already been predicted for UHGV vOTUs using Prodigal-gv and is available in the UHGV metadata.

### Prokka

A fork of Prokka v1.14.5^11^ was written that incorporates an initial stage of ORF prediction using Prodigal-gv v2.11.0 (https://github.com/apcamargo/prodigal-gv)^8^. A first gene calling step is used to infer the genetic code most likely adopted by the genome, then the predicted genetic code is used to perform the translation FASTX::Seq, which we updated to accept code 15 (metacpan.org/pod/FASTX::Seq)^16^. The code for this is available at (github.com/telatin/metaprokka). We included publicly available HMMs of the PHROGs database in our Prokka-gv annotations (http://s3.climb.ac.uk/ADM_share/all_phrogs.hmm.gz)^17^. The fork is installable from Bioconda as ‘metaprokka’.

### Pharokka

Pharokka v1.5.0^12^ was updated to include support for pyrodigal-gv implementing pyrodigal-gv as a gene predictor. This is specified by using ‘-g prodigal-gv’ when running Pharokka. The updated code is available on GitHub (https://github.com/gbouras13/pharokka). Pharokka uses tRNAscan-SE for predicting tRNAs^14^.

### Statistical Analyses and Data Visualisation

To test for significance in differences of results, a simple paired T test was performed in R v4.2.2^18^ and P-values were adjusted using the Benjamini-Hochberg procedure^19^. Figure 1 was produced using ggplot2 v3.4.2^20^.

### Supplementary Results

### Prokka-gv Annotations

For Prokka-gv, the largest differences were observed for sequences predicted to use translation table 15, for which Prokka-gv increased the median gene length (median of per genome medians) from 276 to 396 bp for UHGV sequences (43.5% increase), and from 309 to 483 bp for INPHARED sequences (56.3% increase). This was also reflected in an increase of median coding capacity from 66.6% to 86.7% for UHGV, and from 69.2% to 87.3% for INPHARED. As it is commonly used as a phylogenetic marker for bacteriophages, we investigated how commonly the major capsid protein (MCP) could be identified with and without predicted stop codon reassignment^15^. For sequences predicted to use translation table 15, the MCP could be identified on 382/715 (53.4%) sequences with Prokka and this was marginally increased to 386/715 (53.9%) with Prokka-gv.

When investigating the sequences for which translation table 4 was predicted, a substantial increase was also observed for UHGV sequences, with Prokka-gv increasing median median gene length from 319 to 460 bp (44.2%), resulting in an increase of coding capacity from 78.4% to 91.4%. However, the same was not observed for INPHARED sequences predicted to use translation table 4. These sequences observed a modest increase in median median gene length from 573 to 584 bp (1.8%) for Prokka-gv. Median coding capacity was not increased with Prokka and Prokka-gv both obtaining 86.2%.

